# Loop-mediated Isothermal Amplification (LAMP) assay for reliable detection of *Xanthomonas axonopodis* pv. *vasculorum*

**DOI:** 10.1101/2024.02.07.579270

**Authors:** Mitchell Marabella, Julia Howard, Santosh Bhandari, Sally Do, Maya Montoya-Pimolwatana, Yichen Dou, Shefali Dobhal, Dario Arizala, Stefania Montesinos, Sharon A. Andreason, Francisco Ochoa-Corona, Jon-Paul Bingham, Jenee Odani, Daniel Jenkins, Li Maria Ma, Jacqueline Fletcher, James P. Stack, Mohammad Arif

## Abstract

*Xanthomonas axonopodis* pv. *vasculorum* (Xav), the causative agent of sugarcane gumming disease, represents a significant threat to global sugarcane production due to its systemic and destructive nature. Despite the economic implications, a field-deployable, Xav-specific diagnostic tool has not been developed. This resulted in a loop-mediated isothermal amplification (LAMP) assay targeting the *pelL* gene, unique to Xav strains, as a rapid and precise diagnostic assay. The selection of the *pelL* gene was informed by comprehensive *in silico* analyses of Xav genomes and related *Xanthomonas* species and other close relatives. Validation against the NCBI GenBank database and internally sequenced genomes confirmed the gene’s exclusivity to Xav. Subsequent primers for both endpoint PCR and LAMP assays were designed using the *pelL* gene region. The LAMP assay underwent extensive testing against inclusivity and exclusivity panels. Use of exclusivity panel, comprising 81 strains from related species, other bacterial genera, and host genomes, demonstrated the assay’s specificity with no false positives. The assay exhibited a detection limit of 1 pg, and its effectiveness was unimpeded by crude host lysate (sugarcane). Further validation through multi-device and multi-operator testing underscored the assay’s 100% reproducibility and robustness. Application to infected plant samples resulted in the detection of all infected specimens without any false positives or negatives. This novel LAMP assay is accurate and reliable tool for Xav detection, with promising applications in routine diagnostics, biosecurity measures, microbial forensics, and epidemiological research.

## Introduction

Members of the Gamma proteobacterial genus *Xanthomonas*, composed of 36 validly published species, predominantly infects plants and have caused severe damage to a wide variety of economically important vegetable and field crops and tree plant species^1–3^. Sugarcane (*Saccharum officinarum* L.) is among these plant species affected by *Xanthomonas* species^2,4^*. Xanthomonas axonopodis* pv*. vasculorum* (Xav), a Gram-negative phytopathogenic bacterium, poses a particularly significant threat to the agricultural industry due to its capability to cause gumming diseases in sugarcane, a tropical grass native to Asia. Sugarcane holds a vital position in global sugar production, contributing to around 75% of the industry^5,6^. Gumming disease is a vascular ailment distinguished by its unique symptoms, including distinctive lesions, red discoloration at nodes, gum pockets at growing points, and damage to nodal and internodal tissues^6,7^. These symptoms are followed by chlorosis of new leaves in mature plants, reducing photosynthetic efficiency and consequently decreasing crop yield. Xav demonstrates clear phylogenetic differentiation from other sugarcane-infecting xanthomonads, such as *X. vasicola* pv. *vasculorum*, *X. sacchari*, and *X. albilineans*^2^. Therefore, an accessible, highly specific, and rapid identification method is crucial for reliable diagnosis, efficient disease management, timely intervention, and developing strategies to protect agricultural productivity.

Traditional serological assays, such as the enzyme-linked immunosorbent assay (ELISA), have been used since the 1970s to detect microbial proteins associated with specific pathogens. While genomic data is not required, ELISA is labor-intensive and is often associated with a high incidence of false negatives and false positives, increasing the risk of disease transmission and subsequent crop loss^8^. The advent of PCR has enabled efficient processing, allowing for the precise quantification of amplification products and facilitating the rapid detection of pathogens^9^. Various reliable identification methods, such as endpoint PCR and qPCR assays, have been established for *Xanthomonas* species, offering improved accuracy^10–12^. However, certain limitations have been identified regarding point-of-need applications. Most notably, PCR requires intricate, costly machinery and precise operating conditions^13^.

In contrast, isothermal methods are user-friendly and field-deployable, proving highly advantageous in point-of-need applications^14,15^. Among these isothermal methods, loop-mediated isothermal amplification (LAMP) is particularly favorable for point-of-need assays^16,17^. LAMP assays are widely recognized for their exceptional cost-efficiency and have gained prominence for their utility in various applications^13,18^. Due to its high sensitivity, inhibitor resistance, ease of use, and quick reaction time (under 20 mins), LAMP ensures effectiveness and reliability, even in challenging conditions^19–21^.

LAMP operates by utilizing a DNA polymerase that can displaces strands and synthesize DNA, enabling sequence amplification under isothermal conditions^13,22^. This process relies on four primers comprised two inner primers, the forward inner primer (FIP) and backward inner primer (BIP), and two outer primers, F3 and B3^13^. In the initial stages of LAMP, the inner primer, either FIP or BIP, hybridizes to its target, allowing the polymerase to begin synthesizing the complementary strand. Next, the outer primer, F3 or B3, hybridizes to its target site and displaces the synthesized complementary strand, releasing the strand that serves as a template for the other inner primer. Loop primers, developed by Nagamine and Notomi^22^, accelerate the reaction by hybridization of regions between the binding sites of F3 and FIP, or B3 and BIP, enhancing the formation of the characteristic “dumb-bell” stem-loop structure of the LAMP product with improved selectivity^22,23^. Results of LAMP can be verified by electrophoresis, precipitation of magnesium pyrophosphate byproduct, or SYBR green assay^17,24^. SYBR green assays use a fluorescent intercalating dye present in the solution that binds to double-stranded DNA and can be confirmed visually with the naked eye or under UV light^18^.

The objective of this study was to develop a point-of-need LAMP assay for the specific detection and identification of Xav in infected plant tissues. This objective encompasses the identification of a unique genomic region present in Xav through comparative genomic analyses. These analyses involve *in silico* validation against both public and customized databases, enabling the design of a LAMP primer set. Protocol development includes thorough validation, with extensive exclusivity and inclusivity panels. The developed assay exhibits potential applications in plant disease diagnostics, seed certification, agricultural surveys, farm management, and plant biosecurity.

## Results

### Target Selection and Primer Design

*In silico* analyses identified potential target regions specific to Xav. A region of the *pelL* gene emerged as distinctive to Xav and exhibited conservation across all six strains, thereby establishing its utility for diagnostic assay development. Among the 1,326 nucleotides encoding *pelL*, NCBI BLASTn results revealed that the last 663 base pairs (bp) of the coding region demonstrated discriminating specificity. Primers designed for both PCR and LAMP (Table 1), were subjected to *in silico* validation through BLASTn against an in-house custom database comprising genomes of Xav and closely related species, affirming their high specificity. The location of *pelL* gene/primers is shown in Figure 1.

**Figure 1.**
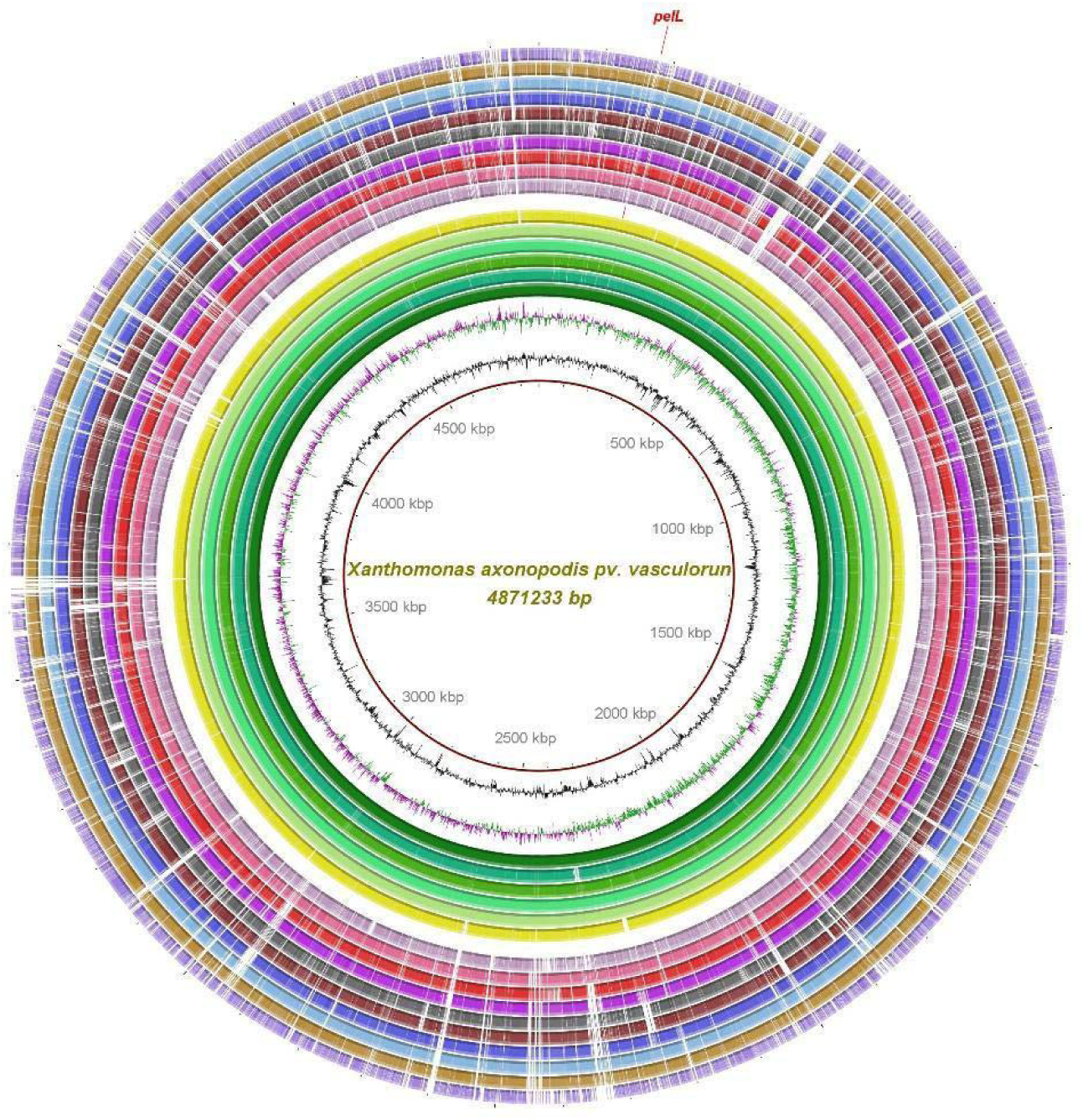
Depicts a ring plot that highlights the unique sequence region employed in developing a loop-mediated isothermal amplification (LAMP) assay for the specific detection of *Xanthomonas axonopodis* pv*. vasculorum*. This circular graphic illustrates multiple genome alignments that include six *X. axonopodis* pv. *vasculorum* genomes (three of which were obtained from our lab) and ten other *Xanthomonas* species that share the same ecological niche or infect the same host, sugarcane. The first three innermost layers of the graphic represent the genome coordinates (in megabase pairs, mbp), the GC content (depicted as a zigzag black line), and the GC skew (shown as a zigzag purple + / green - line) of the reference genome *X. axonopodis* pv. *vasculorum* strain NCPPB 796. Subsequent color-coded rings display the BLASTn pairwise comparisons of various *X. axonopodis* pv. *vasculorum* strains, including NCPPB 796 (NZ_CP053649), NCPPB 900 (NZ_JPHD00000000), CFBP5823 (MDCD00000000), LMG894, LMG901, and LMG903. These are followed by the position of the unique target region conserved across all *X. axonopodis* pv. *vasculorum* strains, which is pointed out and labeled in red. Additionally, the plot includes genomes from other *Xanthomonas* species, such as *X. campestris* pv. *campestris* ATCC 33913 (NC_003902), *X. campestris* pv. *musacearum* NCPPB 4379 (NZ_AKBF00000000), *X. citri* pv. *citri* MN12 (NZ_CP008998), *X. euvesicatoria* pv. *alfalfae* CFBP3836 (NZ_CP072268), *X. oryzae* pv. *oryzae* ICMP3135 (NZ_CP031697), *X. oryzae* pv. *oryzicola* GX01 (NZ_CP043403), *X. phaseoli* pv. *dieffenbachiae* LMG695 (NZ_CP014347), *X. perforans* GEV872 (NZ_CP116305), *X. vasicola* pv. *vasculorum* Xv1601 (NZ_CP025272), and *X. sacchari* DJ16 (NZ_CP121698). This image was generated using the BLAST Ring Image Generator (BRIG) v 0.95^25^.

**Table 1.**
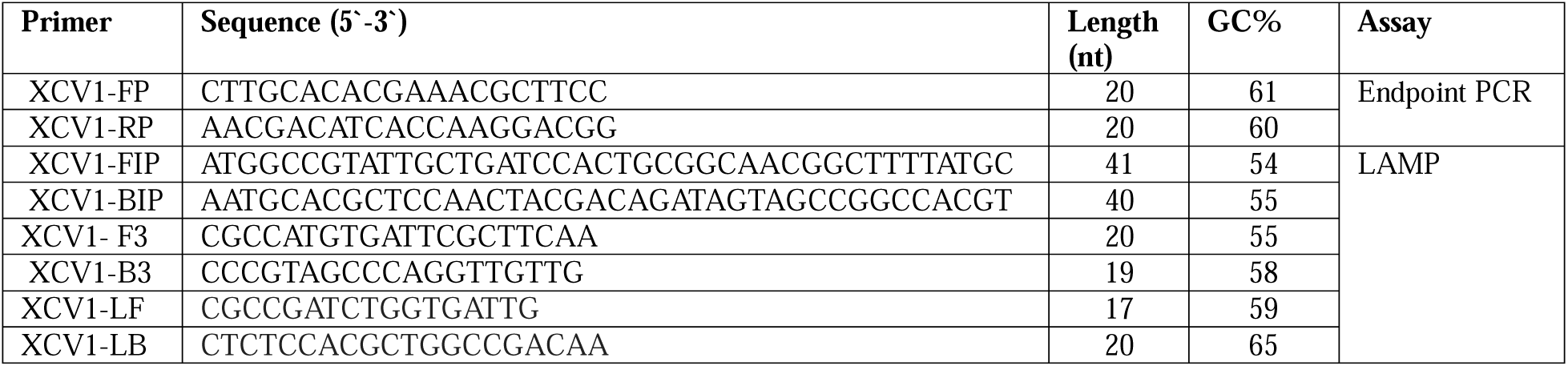
Details of endpoint PCR and loop-mediated isothermal amplification (LAMP) primers designed for specific detection of *Xanthomonas axonopodis* pv. *vasculorum* using the *pelL* gene region.

### End-Point PCR Specificity

The specificity of the endpoint PCR primer set was validated in vitro by testing against several strains included in the inclusivity and exclusivity panels. The inclusivity assay utilized genomic DNA from six strains of Xav (Table 2), while the exclusivity panel comprised three strains of *Stenotrophomonas* and 47 strains of various Xanthomonas species (Table 2). The targeted *pelL* gene was found in all strains of Xav. No off-target amplification of the gene occurred with the strains included in the exclusivity panel. The PCR amplification of the target region using forward and reverse primers XCVI-FP and XCVI-RP yielded a product size of 208 nucleotides, as predicted with Primer3.

**Table 2.**
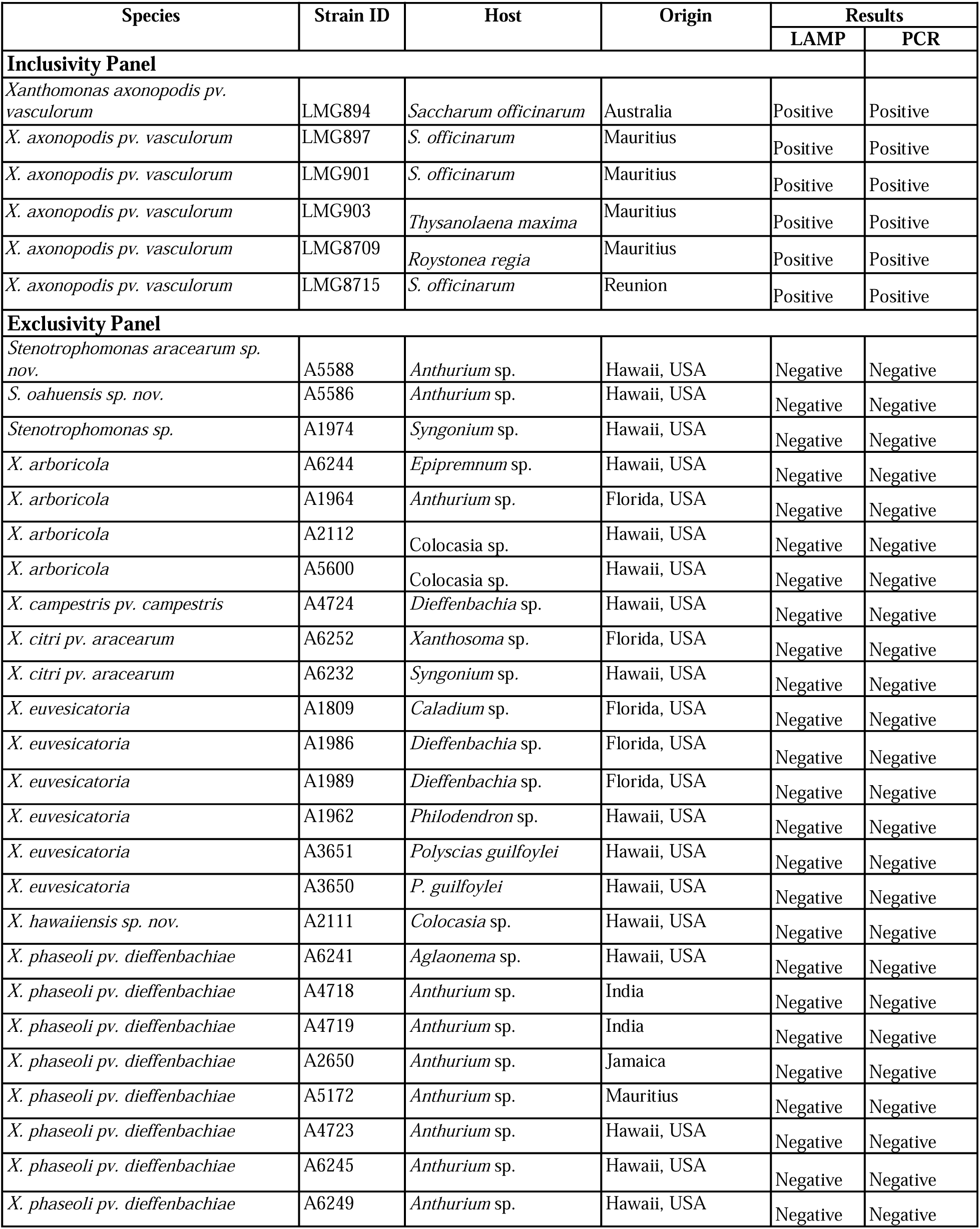
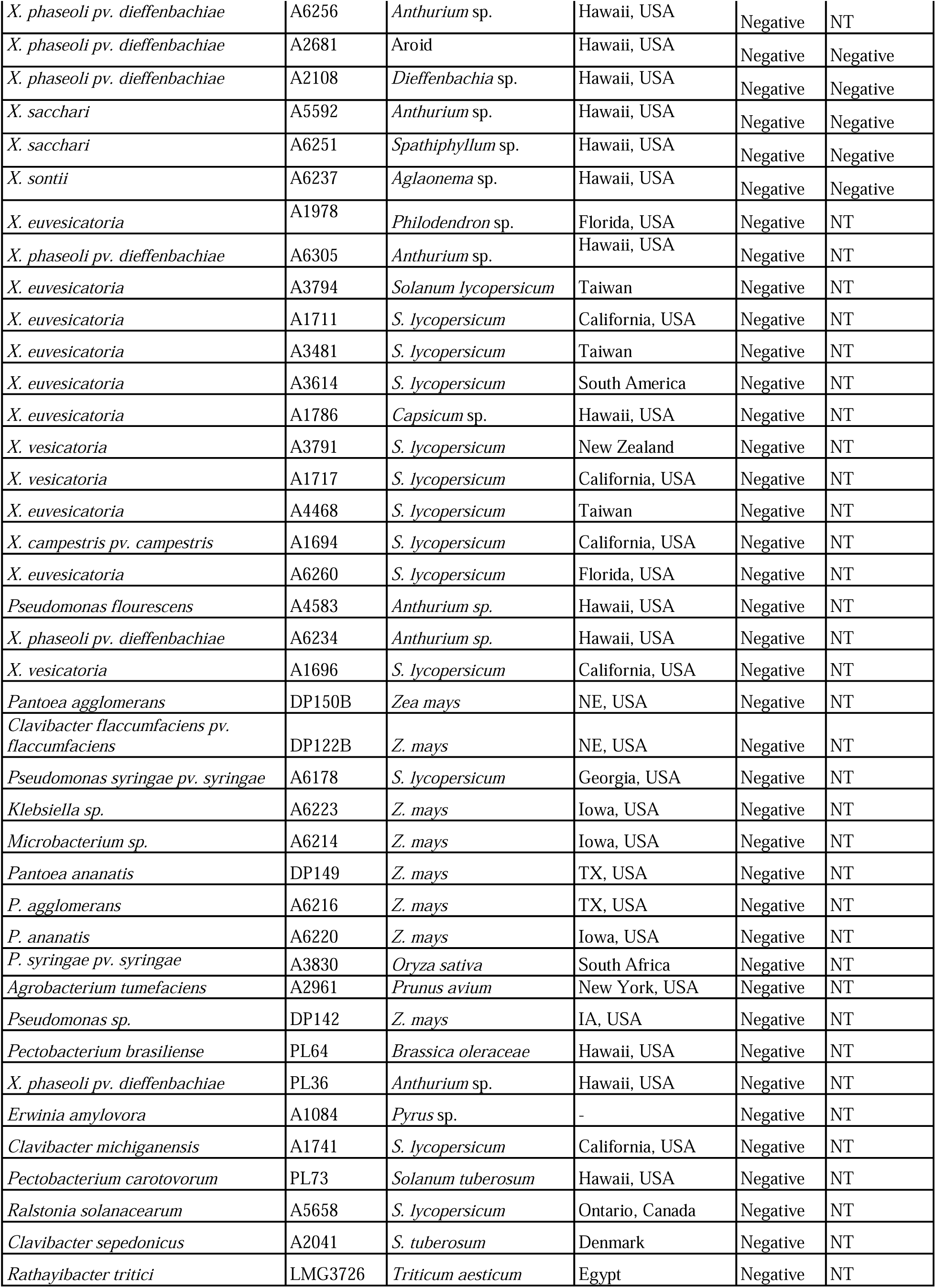
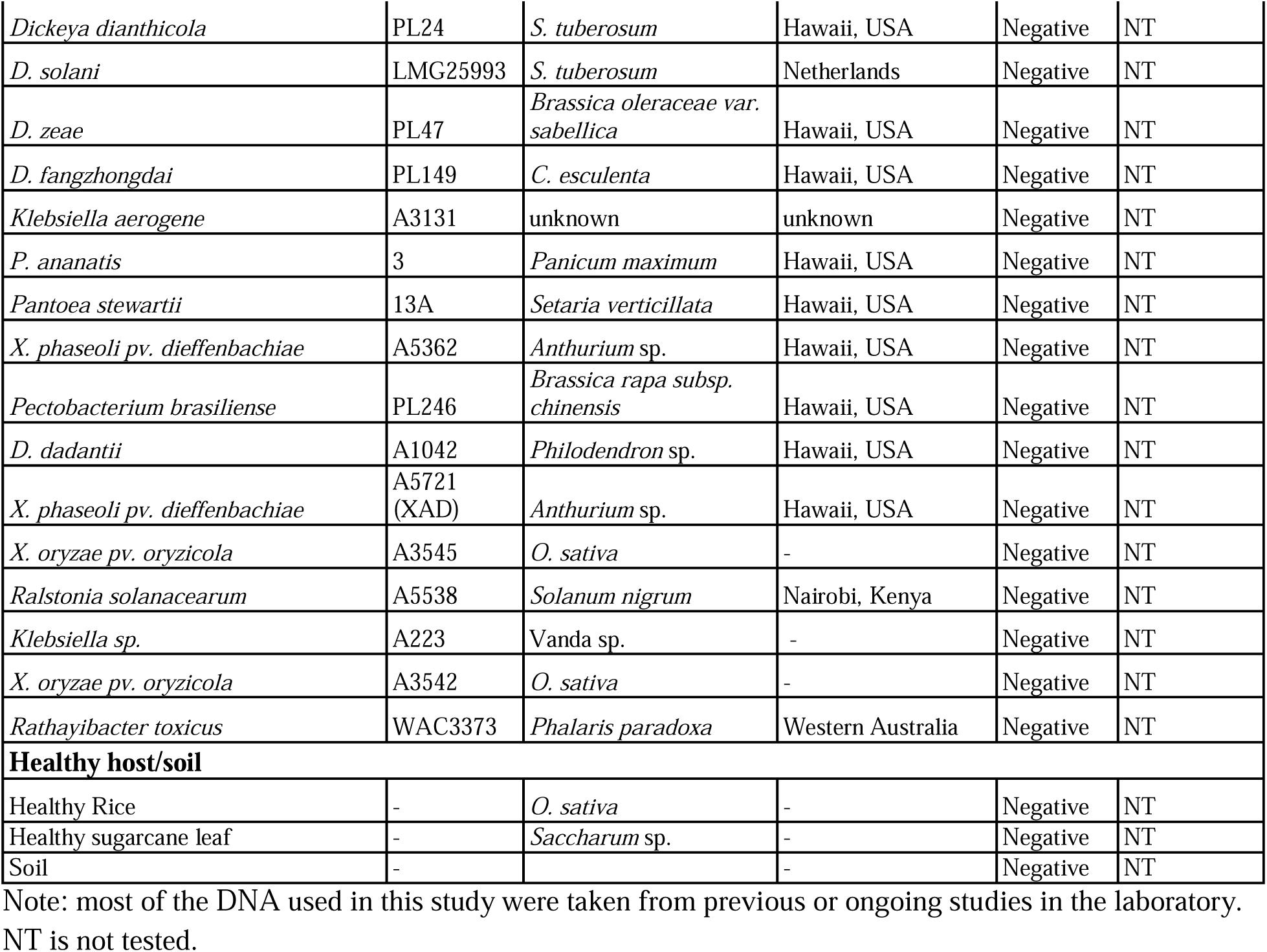
List of strains used in the inclusivity and exclusivity for the validation assay of *Xanthomonas axonopodis* pv*. vasculorum* specific loop mediated isothermal amplification (LAMP) and endpoint PCR.

### LAMP Assay Specificity

We employed an extensive panel of strains to demonstrate the specificity of the LAMP primers designed using the *pelL* gene region (Tables 1 and 2). The inclusivity panel comprised six strains of Xav, while the exclusivity panel included the remaining strains (Table 2). The exclusivity panel was selected based on phylogenetic relationships, common niche, and geographical locations. Following amplification, the assay detected all six Xav strains from the inclusivity panel, and no false positives were obtained from any strains in the exclusivity panel. Quantification of isothermal cycling data produced sigmoidal curves indicating a positive result surpassing a threshold of 0.2 normalized fluorescence (Figure 2A). Positive samples displayed Ct values ranging from 8.76 (LMG8715) to 14.45 (LMG894) (Figure 2A). Amplified DNA was confirmed colorimetrically by the appearance of fluorescent green (Figure 2B), and also detectable through exposure to UV light fluorescence (Figure 2C). SYBR Green detection was concordant with the results of sigmoidal amplification curves.

**Figure 2.**
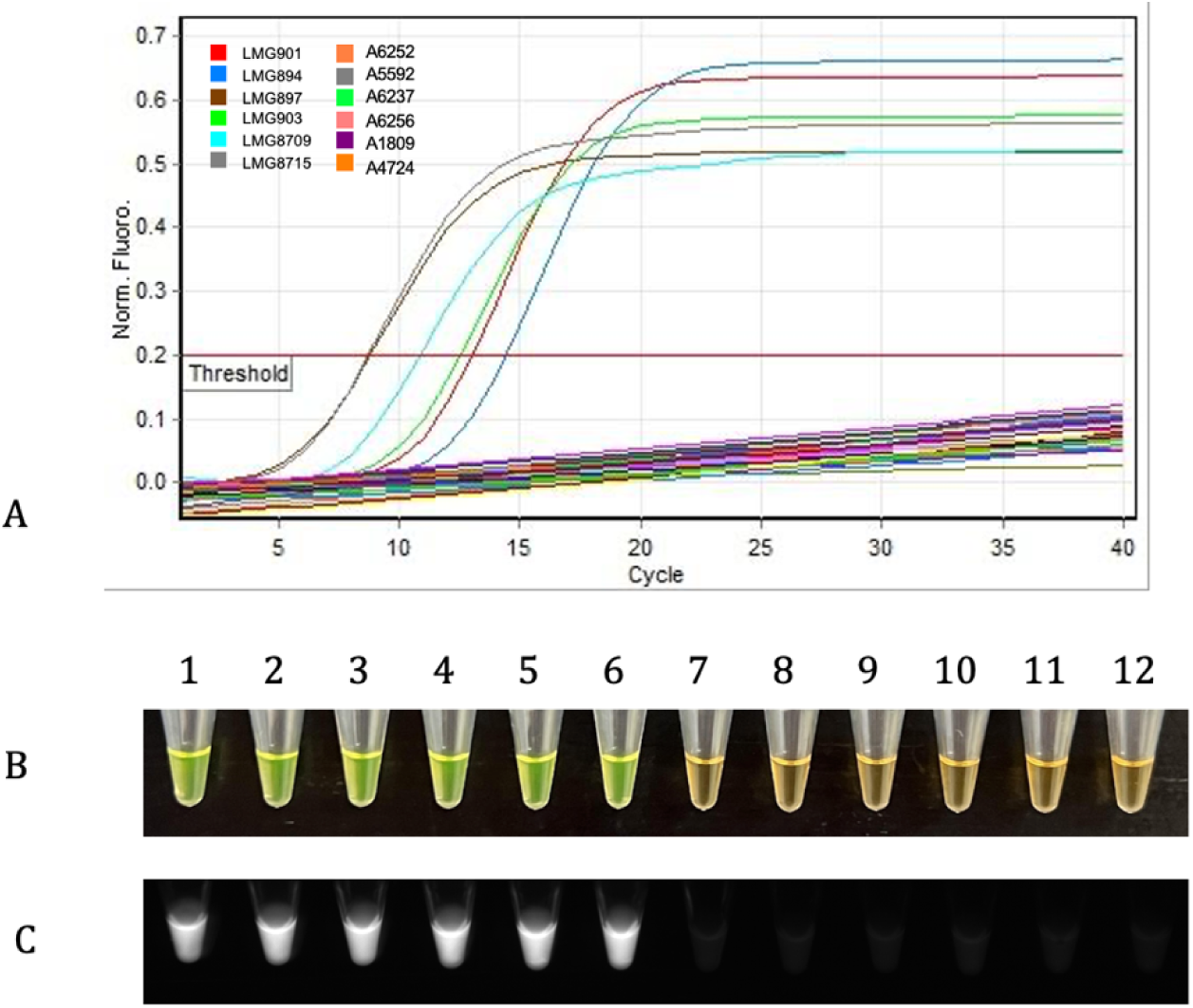
Specificity validation of the loop-mediated isothermal amplification (LAMP) assay designed for specific amplification of *Xanthomonas axonopodis* pv. *vasculorum* was conducted with 87 bacterial strains, as detailed in the inclusivity and exclusivity panels in Table 2. (A) Curves surpassing the threshold of 0.2 normal fluorescence indicate a positive result. All strains from the inclusivity panel were positive, and all strains from the exclusivity panel tested negative. (B) SYBR Green I dye was added to each tube, with Xav strains in tubes 1-6 (LMG901, LMG894, LMG897, LMG903, LMG8709, LMG8715) and exclusivity strains in tubes 7-12 (A6252 - *X. citri* pv. *aracearum*, A5592 - *X. sacchari*, A6237 - *X. sonti*, A6256 - *X. phaseoli* pv. *dieffenbachiae*, A1809 - *X. euvesicatoria*, A4724 - *X. campestris* pv*. campestris*). Positive results changed to green, while negative results remained orange. (C) Positive SYBR Green binding was verified by fluorescence emission detection under UV light.

### Limit of detection

The detection limit of the LAMP assay was determined using four independent sensitivity assays: 1) artificial positive control, 2) purified genomic DNA, 3) purified genomic DNA spiked with crude lysate from host leaves, and 4) purified genomic DNA spiked with crude lysate from host stems. For these assays, tenfold serially diluted genomic DNA and plasmid DNA (only for artificial positive control) were used.

The LAMP assay was sensitive, detecting purified genomic DNA, with and without adding host crude lysate, down to 1 pg per reaction (Figure 3). The introduction of crude host lysate extracted from the leaves and sets (stems) of sugarcane showed no adverse effect on detection, confirming the applicability of this assay for in-field detection without purifying high-quality genomic DNA. Validation across multiple detection chemistries, including SYBR Green fluorescence, UV, and gel electrophoresis, revealed consistent outcomes, with no discrepancies among the respective results.

**Figure 3.**
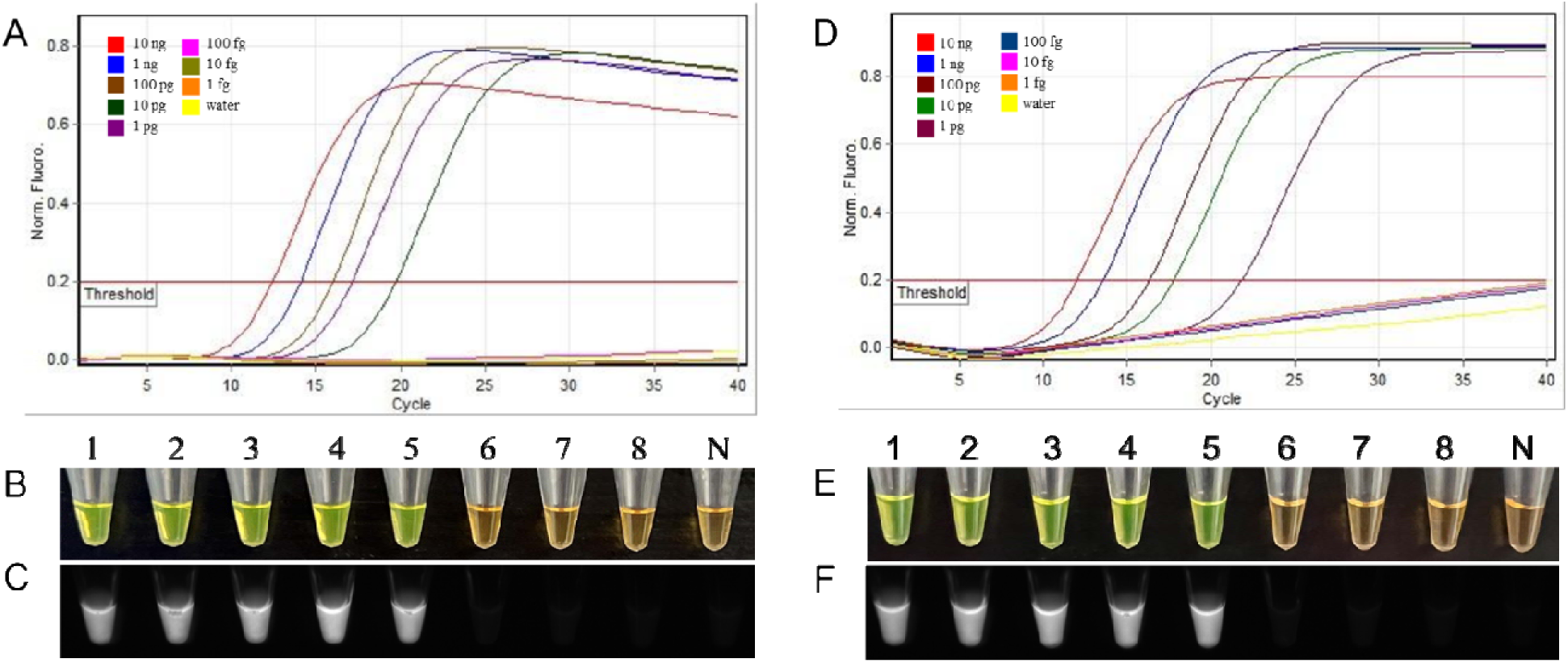
Limit of detection (LoD) determination of the loop-mediated isothermal amplification (LAMP) assay using serially diluted purified genomic DNA and genomic DNA spiked with crude host lysate (sugarcane stem). The purified genomic DNA of strain LMG901 was tenfold serially diluted (from 10 ng to 1 fg) and added to the LAMP reaction tubes (1–8). For the spiked assay, 5 µl of crude host lysate was added in each tube containing 10-fold serially diluted genomic DNA. Tube N: Non-template control (NTC) water. Figures 3A to 3C and 3D to 3F represent results from sensitivity and spiked sensitivity assays, respectively. A standardized sigmoidal curve (A and D); SYBR Green verification of results performed by adding SYBR Green I dye to the amplified reaction tubes—a color change from orange to bright green indicates positive amplification (B and E); UV fluorescence confirmation of SYBR Green added tubes - positive amplification emit fluorescence (C and F). The LAMP assay detected amounts a low as 1 pg with both sensitivity and spiked sensitivity assays.

The limit of detection with a positive control was performed using purified plasmid DNA containing the target region of the *pelL* gene of Xav. LAMP primers F3 and B3 were used to amplify the *pelL* gene region. The amplified region was inserted into DH5-alpha *E. coli*, and the isolated plasmid DNA was used for the sensitivity assay. The limit of detection using plasmid DNA was 1 fg.

### Multi-operator Blind Validation Test

The viability and robustness of the LAMP assay in a point-of-need setting were tested through a blind multi-operator test. Two independent operators examined ten unknown samples containing Xav and other closely related species. Each operator conducted the assay independently, and both correctly identified three Xav strains and seven non-Xav samples, including a non-template control, with 100% accuracy (Figure 4).

**Figure 4.**
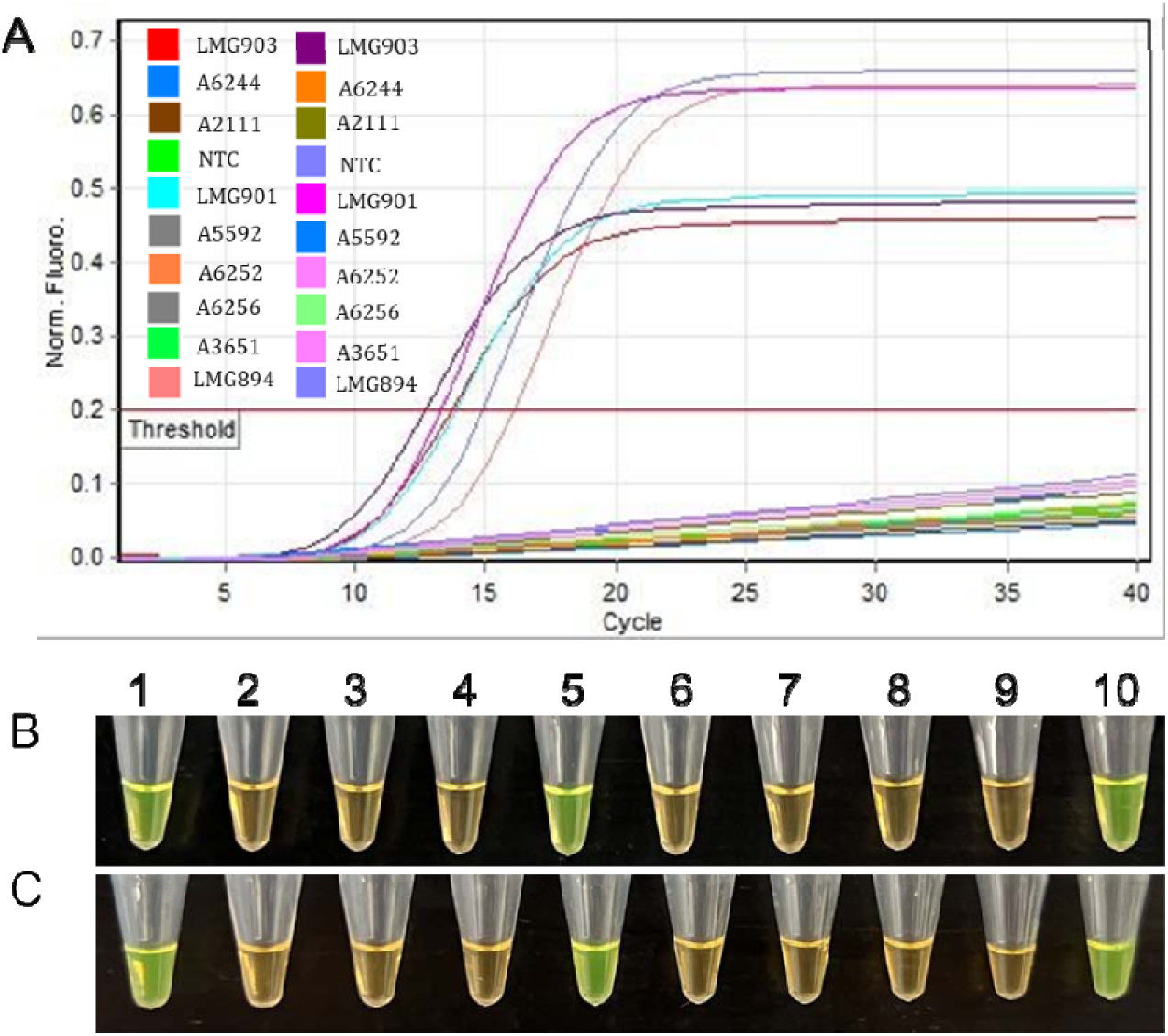
Multi-operator blind tests were conducted to assess the reproducibility and robustness of the loop-mediated isothermal amplification (LAMP) assay, designed for the specific detection of *Xanthomonas axonopodis* pv. *vasculorum* (Xav). Ten representative samples from inclusivity and exclusivity panels were used in each blind test. (A) Standardized sigmoidal curve, incorporating results from both operators. (B) and (C) results from operator 1 and operator 2, respectively, after the addition of SYBR Green dye I. The color change from orange to bright green indicates positive amplification. Tubes 1, 5, and 10 were correctly identified as Xav (strains LMG903, LMG 901, LMG894). Non-template control (NTC; tube 4).

### Multi-Device Validation

Multi-device validation of the developed assay was conducted using the T100 thermal cycler (Bio-Rad, Hercules, CA), Rotor-Gene Q real-time PCR system (Qiagen, Germantown, MD), and a digital heat block (VWR) to ensure its reproducibility. The validation involved testing ten samples, including five strains of Xav: LMG901, LMG894, LMG897, LMG903, and LMG8709. The other five samples comprised *X. campestris* pv. *campestris* (A4724), *X. sacchari* (A5592), *X. citri* pv. *aracearum* (A6252), *X. sontii* (A6237), and a non-template control. The results obtained from each device were concordant, with no false positives or negatives, thereby confirming the high reproducibility and reliability of the assay (Figure 5).

**Figure 5.**
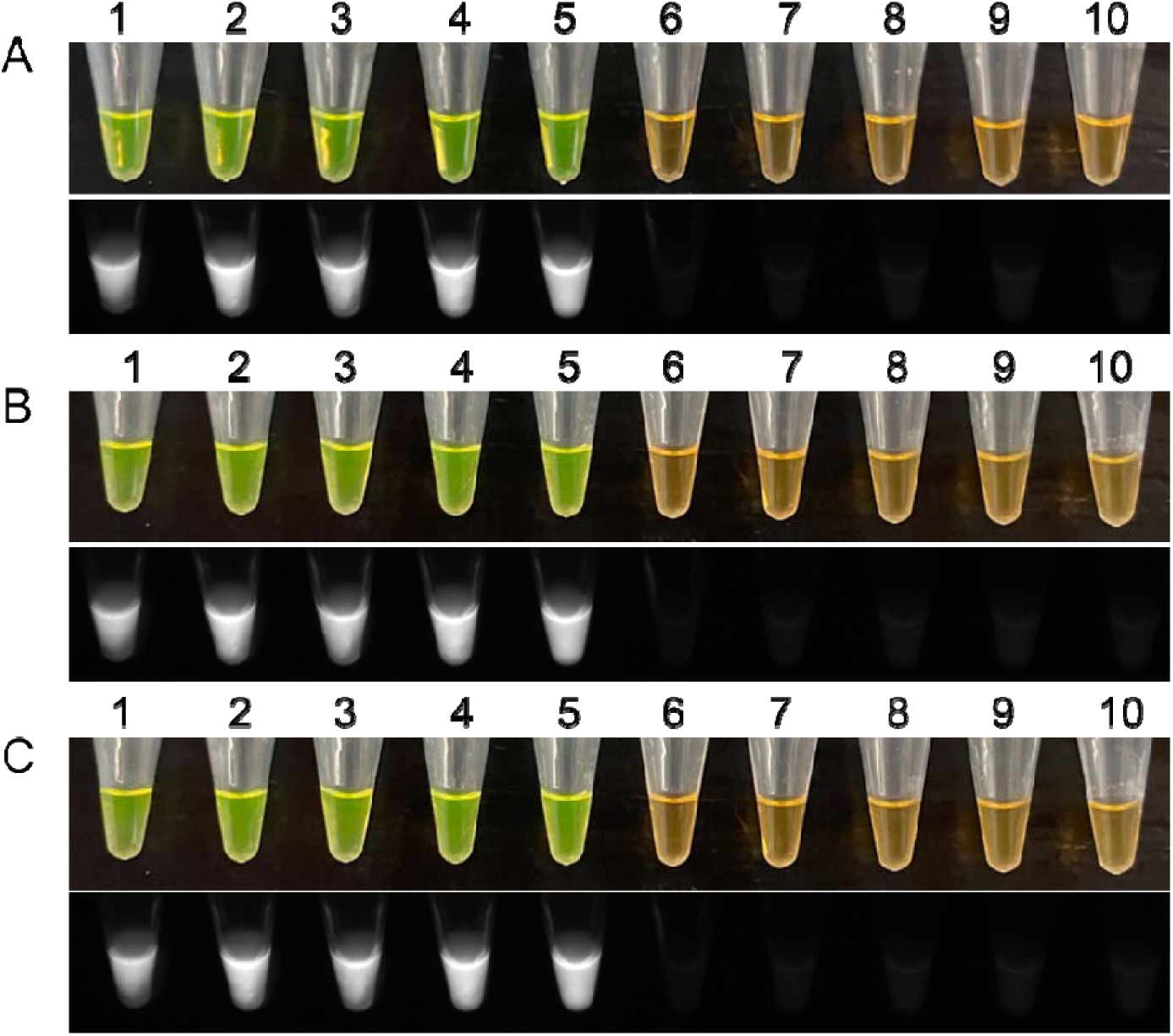
Multi-device validation of *Xanthomonas axonopodis* pv. *vasculorum* was conducted using three different instruments: (A) Rotor-Gene Q real-time PCR (Qiagen), (B) T100 thermal cycler (Bio-Rad), and (C) dry bath (VWR). Three independent operators performed the assays on the aforementioned platforms, maintaining a consistent temperature of 65 °C for 20 minutes. The positive samples comprised cell lysate directly from Xav colonies (LMG901, LMG894, LMG897, LMG903, LMG8709). The change to green color after addition of SYBR Green dye and subsequent fluorescence under UV light indicated positive amplification. Five non-target samples were included: *X. campestris* pv. *campestris* (A4724), *X. sacchari* (A5592), *X. citri* pv. *aracearum* (A6252), *X. sontii* (A6237), and a non-template control. After addition of SYBR Green dye, these samples remained their orange and showed no fluorescence under UV light, indicating a negative reaction.

### Detection from Artificially Inoculated Samples

To evaluate the developed LAMP assay for the application of *on-site* detection, we utilized six strains of Xav (LMG894, LMG897, LMG901, LMG903, LMG8709, and LMG8715) for inoculation of sugarcane slices. Additionally, we used other *Xanthomonas* species for inoculation, including *X. campestris* pv. *campestris* (A4724), *X. hawaiiensis* sp. nov. (A2111), *X. citri* pv. *aracearum* (A6252), *X. sacchari* (A5592), and *X. sontii* (A6237)^26^. Two control groups were included: one with healthy sugarcane and another with a non-template control. The LAMP assay successfully detected Xav in the infected sugarcane plant slices.

## Discussion

The genus *Xanthomonas* poses a significant threat to many important crops, with the potential for substantial economic losses^3,27,28^. Given the risks associated with this genus, various assays suitable for point-of-need detection of many *Xanthomonas* spp. have been developed^29–32^. Despite the threat posed by gumming disease in sugarcane, no such assay is currently available for Xav. The field-deployable LAMP is a widely used and well-established method for pathogen detection and diagnosis due to its cost-effectiveness, high sensitivity, and ease of use^16,29,33^. Consequently, we have developed a robust LAMP assay for the specific detection of the sugarcane systemic pathogen Xav.

Target selection is of paramount importance to develop a reliable, robust, and specific diagnostic assay^14,34–36^. The genomes of Xav were compared against closely related species and other bacterial genera to identify a Xav-specific genomic region. This comparative genomics analysi identified a unique region in a portion of the pectate lyase (*pelL*) gene conserved exclusively in all Xav strains. Furthermore, the *pelL* gene region was chosen due to its predicted conserved function. The endpoint PCR and LAMP primers designed using this *pelL* gene region were specific to Xav strains only.

In the case of multiple diseases, caused by multiple pathogens (especially if they are in closely related taxa), affecting a common host, diagnostic techniques should be optimized to be highly specific, avoiding cross-reactivity with other species and limiting false positives for proper disease management^37,38^. The LAMP assay developed for other diseases were observed to be both highly selective and sensitive^29,33,39^. The specificity of the Xav-specific LAMP assay was verified by inclusivity/exclusivity assays containing multiple strains of Xav and 81 strains of non-target bacterial species.

High sensitivity was confirmed through four Limit of Detection (LoD) assays. Firstly, an artificial positive control (APC) was generated from the target sequence, *pelL* gene, using primers F3 and B3 and subsequently cloned. The resulting plasmid created from this process provides users with a reliable, and readily available positive control for the Xav-specific LAMP assay. The APC facilitated further comparison and validation of an accurate LoD without uncertainty related to inhibitor presence or nucleotide extraction procedures^34^. As plant biosecurity measures often restrict the import of many pathogens, the development of an APC enables regions that cannot obtain a positive control for Xav to instead utilize the non-infectious plasmid for assay development and validation, thereby streamlining the assay adaptation process in different labs nationally and internationally^34,40^. The LoD was determined to be 1 fg with plasmid DNA. In a prior study by Arif *et al*.^14^, it was observed that the LAMP assay is relatively less susceptible to plant inhibitors than PCR, however not as effective as the RPA assay. The LoD was verified using Xav genomic DNA and two spiked assays involving different sugarcane tissues (leaf and culm). These assays were conducted to confirm that various infected tissues would not result in false negatives due to inhibitors present in different host tissues. The LAMP assay detected Xav down to 1 pg in all LoD assays, with no signs of inhibition, confirming the robustness and field applicability of the developed assay.

Reproducibility is a fundamental attribute of a diagnostic assay^34^. Consequently, we performed comprehensive assessments to evaluate the LAMP assay’s performance. The evaluation of the LAMP assay involving multiple operators demonstrated consistent and accurate pathogen detection with no discrepancy (Figure 4). The assay exhibited consistent performance across various operating devices, showcasing robustness and versatility among diverse amplification platforms (Figure 5). The assay displayed resilience against plant material inhibitors and maintained accurate diagnosis when tested using crude lysate from artificially infected sugarcane material (Figure 6). These observations suggest that the assay does not require highly purified DNA and can be performed using simple preparation methods^19,30^. Collectively, the assessment highlights the reliability of the developed LAMP assay. Its simplistic protocol makes it suitable for implementation across various laboratories and fields, eliminating the necessity for specialized methods.

**Figure 6.**
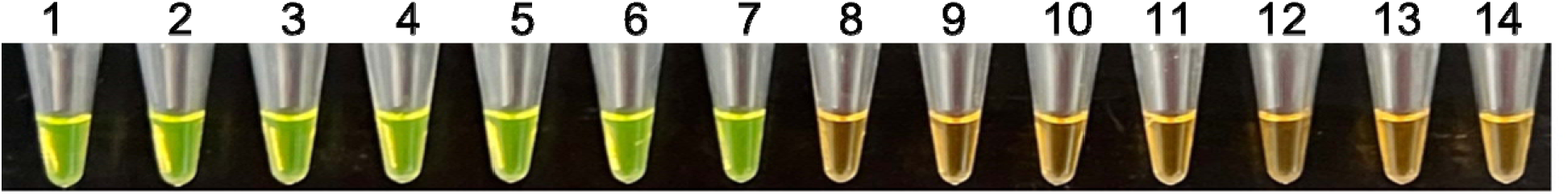
Detection of *Xanthomonas axonopodis* pv. *vasculorum* from infected sugarcane plant slices. LAMP products were visualized after addition of SYBR Green I dye—green color represents positive amplification. Tube 1 is a positive control (LMG901); tubes 2-7 are sugarcane plant slice samples infected with six different Xav strains: LMG894, LMG897, LMG901, LMG903, LMG8709, and LMG8715; tubes 8-12 are sugarcane slices inoculated with other Xanthomonas species: *Xanthomonas campestris* pv. *campestris* (A4724)*, Xanthomonas hawaiiensis* sp. nov. (A2111), *Xanthomonas citri* pv. *aracearum* (A6252)*, Xanthomonas sacchari* (A5592)*, Xanthomonas sontii* (A6237); tube 13 contains healthy sugarcane, and tube 14 is non-template control (NTC – sterile water).

In conclusion, we have developed a Xav-specific LAMP assay using a highly conserved and unique genomic region within the Xav strains. It is robust specific, detecting the target genomic region down to 1 pg. The assay can effectively be deployed in disease diagnosis, farm management, quarantine, border protection, epidemiology, and planting material certification under both laboratory and field conditions.

## Materials and Methods

Any plant and plant materials used in this research comply with international, national, and institutional guidelines.

### Bacterial Isolates Used for DNA Extraction

The current study incorporated 87 bacterial strains from various hosts and geographic locations to represent the six strains of Xav and other taxonomically related species. The inclusivity panel consisted of 6 Xav strains (procured from BCCM/LMG; Bacteria Collection, Belgium), while the exclusivity panel consisted of 81 bacterial strains (various species of *Xanthomonas, Stenotrophomonas, Pantoea, Pectobacterium, Erwinia, Clavibacter, Dickeya*, and *Klebsiella*) from the Pacific Bacterial Collection and Phytobacteriology culture collection (The University of Hawaii at Manoa) and other sources (Table 2). The bacterial strains used in the exclusivity panel were removed from storage (−80 °C) and plated onto various media, including nutrient agar (NA) (BD, Becton Dickinson) glucose NAG media (NA medium supplemented with 0.4% glucose), dextrose peptone agar (DPA) media (10 g 1^-1^ peptone, 5 g 1^-1^ dextrose, 17 g 1^-1^ agar), and 2,3,5-tetrazolium chloride (TZC) media (10 g 1^-1^ peptone, 5 g 1^-1^ dextrose, 17 g 1^-1^ agar, and 0.001% TZC).

The six Xav strains included in the inclusivity panel (strains LMG894, LMG897, LMG901, LMG903, LMG8709, and LMG8715) were revived from lyophilized tubes. Each tube was aseptically broken inside a biosafety cabinet II and then resuspended in 200 µl of dextrose peptone broth and plated on M009 plates (10 g glucose, 5 g yeast extract, 30 g CaCO_3_, 15 g agar, pH adjusted to 7.0 and final volume to 1L) and incubated at 28 °C for 48-72 hours. Single colonies were picked and re-streaked onto another M009 plate. Approximately one-fourth loopful (10 µl) of pure bacterial cells from M009 plates was suspended in phosphate-buffered saline (PBS) contained in 1.5 ml centrifuge tubes, 200 µl of alkaline lysis buffer was added, and genomic DNA was extracted using the Dneasy Blood and Tissue Kit (Qiagen). The same kit was used to isolate DNA from other strains. The DNA of certain strains included in the study were from other studies (S. Dobhal, S. Chuang and M. Arif; unpublished information). DNA of *Xanthomonas oryzae* pv. *oryzae*, *Xanthomonas oryzae* pv. *oryzicola* and *Ralstonia solanacaerum* R3B2 were used from the previous study (Dobhal et al., unpublished information)

### Target Gene Selection

The genomes of three Xav strains, NCPPB796 (NZ_CP053649), CFBP5823 (MCDC00000000), and NCPPB900 (NZ_JPHD00000000), and genomes of other closely related species, were downloaded from the NCBI GenBank genome database. Fragments of strain NCPPB900 were concatenated for alignment, and Geneious Prime 2023.2.1 was employed to identify a unique gene region shared between the Xav strains. All genomes were aligned using Mauve^41^. Gene regions unique to Xav were verified utilizing the nucleotide BLASTn tool. The gene region Pectate lyase (*pelL*) was consistently present across all available genomes, including three available in our lab and three from NCBI GenBank (NCPPB796, CFBP5823, and NCPPB900).

### Primer Specificity with Endpoint PCR

The unique *pelL* region was utilized to design end-point PCR and LAMP primers. End-point PCR primers were designed using Primer3 v4.1.0 (https://bioinfo.ut.ee/primer3-0.4.0), following the procedures outlined by Arif and Ochoa-Corona^42^. The specificity of the designed primers was validated *in silico* using the NCBI database, and a local custom database was created using Geneious Prime. Both primers demonstrated high specificity for Xav.

Each 20 µl endpoint PCR reaction contained 10 µl of Gotaq Green Master Mix (2X) (Promega, Madison, WI), 1 µl (5 µM) of each forward and reverse primer, 7 µl of nucleus-free water, and 1 µl of DNA template. PCR amplification was carried out in a T100 thermal cycler (Bio-Rad) following a thermal cycling protocol including an initial denaturation step at 95 °C for 3 minutes, followed by 35 cycles of denaturation at 95 °C for 30 seconds, annealing at 57 °C for 30 seconds, extension at 72 °C for 30 seconds, and a final extension at 72 °C for 3 minutes.

For electrophoresis, 10 μl of the PCR product was loaded onto a 1.5% agarose gel stained with ethidium bromide (Invitrogen, Carlsbad, CA). The electrophoresis was conducted at 100 V for 40 minutes. PCR results were visualized using UV light under a gel documentation system (Bio-Rad Gel Doc-XR+; Bio-Rad).

### LAMP Assay

LAMP primers were designed using PrimerExplorer V5 (https://primerexplorer.jp/e/). The primers comprised the forward inner primer (XCV1-FIP), backward inner primer (XCV1-BIP), forward outer primer (XCV1-F3), backward outer primer (XCV1-B3), loop forward primer (XCV1-LF), and loop backward primer (XCV1-LB). Two additional primers were designed manually to expedite the reaction. All primers are detailed in Table 1. *In silico* validation was performed following the aforementioned procedure, demonstrating specificity with Xav.

Each 25 µl LAMP reaction contained 15 µl of Isothermal Master Mix (Optigene; ISO-001), 2.0 µl of the LAMP Primer Mix, 7.0 µl of nucleus-free (NF) water, and 1 µl of template DNA. The LAMP primer mix comprised LAMP primers XCV1-FIP (1.6 µM), XCV1-BIP (1.6 µM), XCV1-F3 (0.2 µM), XCV1-B3 (0.2 µM), XCV1-LF (0.4 µM), and XCV1-LB (0.4 µM). The positive control included the Xav strain LMG901, while the non-template control (NTC) consisted of nuclease-free water. For detection, the reactions were carried out at 65 °C for 20 minutes using the Rotor-Gene Q (Qiagen). Melt curve analysis was conducted from 99 °C to 80 °C, increasing at 0.2 °C per second, using Rotor-Gene Q software v.2.3.1 (Built 49). During amplification, samples with Ct value above the threshold of 0.2 were considered positive.

Amplification of the target region was confirmed by adding 3 µl of SYBR Green I solution (Life Technologies Corporation, Eugene, OR) to each reaction. Positive results were manifested by a green color, indicating successful binding of the dye to the amplified DNA, whereas negative samples retained an orange color. SYBR Green binding was further validated through fluorescence detection in a gel doc system under UV light.

### Artificial Positive Control (APC)

An artificial positive control was generated by amplifying the target Xav DNA with F3 and B3 primers. The amplified product was ligated into pJET 1.2/blunt vector using a CloneJET PCR cloning kit (Thermo Fischer Scientific Inc., Worcester, MA) and the ligated product was transformed into *E. coli* DH5 alpha by heat shock. Transformed cells were plated on LB media containing carbenicillin (100 µg/ml) to confirm the successful integration of DNA into the plasmid. The plates were incubated at 37 °C for 24 hours. Subsequently, 17 bacterial colonies were randomly selected and confirmed carrying the target gene region using the aforementioned primers. The colonies were further inoculated in LB broth supplemented with carbenicillin and incubated overnight in a shaker at 120 rpm at 37°C. The plasmid DNA was extracted using QIAprep Spin Miniprep Kit (Qiagen) and the plasmid DNA was verified for the cloned PCR fragment with Sanger sequencing using the pJET_F and pJET_R primer set.

### Limit of Detection

Four assays were conducted to determine the limit of detection (LoD) of the LAMP assay. The first LoD was confirmed using tenfold serially diluted APC (10 ng to 1 fg). The LoD with genomic DNA was assessed by tenfold serially diluted genomic DNA of Xav strain LMG901 (pathotype strain), ranging from 10 ng to 1 fg. Also, two independent assays spiked with sugarcane leaf tissues and cane were performed. In each spiked assay, 5 µl of crude lysate from healthy sugarcane leaf tissue or cane was added to each tube containing tenfold serially diluted genomic DNA from 10 ng to 1 fg. The crude lysate (unpurified DNA) was extracted using a plant material lysis kit (Optigene, West Sussex). An NTC was included in each reaction.

### Multi-operator Blind Test

For additional validation of the LAMP assay, two independent operators conducted blinded LAMP assays. Ten blinded samples were prepared, encompassing strains from both inclusivity and exclusivity panels, along with a no-template control (NTC). The operators adhered to the identical protocol outlined above. The outcomes were corroborated through Rotor-Gene Q melting curve analysis, and SYBR Green I was added for further validation.

### Multi-device Test

Three independent operators conducted a multi-device assessment utilizing three distinct platforms, each maintained at a temperature of 65 °C for 20 minutes. The three incubation platforms employed in this study were the Bio-Rad T100 thermal cycler, Qiagen Rotor-Gene Q, and a dry bath system. Positive samples consisted of crude lysate extracted from Xav colonies (LMG901, LMG894, LMG897, LMG903, LMG9708), while four strains of *Xanthomonas* spp. Were employed as negative samples (A4724, A5592, A6252, A6237), with one sample acting as the NTC. For this assay, the strains were cultured on M009/NAG media at 28 °C for 2 days. The colonies were picked from the media plate, suspended in 50 μL of nuclease-free water, and subjected to heating at 95 °C for 10 minutes in a thermocycler to obtain crude cell lysate. All tubes were placed on ice, followed by a brief spin for 1 minute, and the supernatant was removed and transferred to another tube. One microliter of crude cell lysate was used directly in the LAMP reaction.

### Detection from Artificially Inoculated Plant Samples

To evaluate LAMP assay’s accuracy for on-site detection, six Xav strains (LMG894, LMG897, LMG901, LMG903, LMG8709, and LMG8715) were used for sugarcane plant slice inoculation. Prior to inoculation, healthy sugarcane stalks were washed with 10% sodium hypochlorite solution for 5 minutes, rinsed three times with sterile water, and sliced. Each slice was inoculated with 10 μl of 10^8^ CFU/ml of the Xav strains (Table 2, inclusivity panel) or other *Xanthomonas* spp.: *X. campestris* pv. *campestris* (A4724), *X. hawaiiensis* sp. nov. (A2111), *X. citri* pv. *aracearum* (A6252), *X. sacchari* (A5592), and *X. sontii* (A6237)^26^. The plates were incubated for 2 days at 28 °C. To evaluate the field applicability of LAMP assay, uninfected and infected sugarcane tissues were processed using the Plant Material Lysis Kit following the manufacturer’s protocol, and 5 μL of crude lysate was directly used in the LAMP reaction. Each sample was amplified using the Rotor-Gene Q at 65 °C for 20 minutes.

## Data Availability

The genomes were retrieved from the NCBI GenBank database and are available under the following accession numbers:

*X. axonopodis* pv. *vasculorum* NCPPB796 (NZ_CP053649) https://www.ncbi.nlm.nih.gov/datasets/genome/GCF_013177355.1/;

*X. axonopodis* pv. *vasculorum* CFBP5823 (MCDC00000000) https://www.ncbi.nlm.nih.gov/datasets/genome/GCF_002939725.1/;

*X. axonopodis* pv. *vasculorum* NCPPB900 (NZ_JPHD00000000) https://www.ncbi.nlm.nih.gov/datasets/taxonomy/325777/;

*X. campestris* pv. *campestris* ATCC 33913 (NC_003902) https://www.ncbi.nlm.nih.gov/datasets/genome/GCF_000007145.1/;

*X. campestris* pv. *musacearum* NCPPB 4379 (NZ_CP034655) https://www.ncbi.nlm.nih.gov/datasets/genome/GCF_000277895.2/;

*X. citri* pv. *citri* MN12 (NZ_CP008998) https://www.ncbi.nlm.nih.gov/datasets/genome/GCF_000961215.1/;

*X. euvesicatoria* pv. *alfalfae* CFBP 3836 (NZ_CP072268) https://www.ncbi.nlm.nih.gov/datasets/genome/GCF_017724035.1/;

*X. oryzae* pv. *oryzae* ICMP3135 (NZ_CP031697) https://www.ncbi.nlm.nih.gov/datasets/genome/GCF_004136375.1/;

*X. oryzae* pv. *oryzicola* GX01 (NZ_CP043403) https://www.ncbi.nlm.nih.gov/datasets/genome/GCF_008370835.2/;

*X. phaseoli* pv. *dieffenbachiae* LMG 695 (NZ_CP014347) https://www.ncbi.nlm.nih.gov/datasets/genome/GCF_001564415.1/;

*X. perforans* GEV872 (NZ_CP116305) https://www.ncbi.nlm.nih.gov/datasets/genome/GCF_028010245.1/;

*X. vasicola* pv. *vasculorum* Xv1601 (NZ_CP025272) https://www.ncbi.nlm.nih.gov/datasets/genome/GCF_003949975.1/;

*X. sacchari* DJ16 (NZ_CP121698) https://www.ncbi.nlm.nih.gov/datasets/genome/GCF_029761895.1/.

## Acknowledgements

This work was supported by the USDA-ARS Agreement no. 58-2040-9-011, Systems Approaches to Improve Production and Quality of Specialty Crops Grown in the U.S. Pacific Basin; sub-project: Genome Informed Next Generation Detection Protocols for Pests and Pathogens of Specialty Crops in Hawaii. This work is also supported by USDA-NIFA Award No. 2023-70440-40179, the Barry and Barbara Brennan Endowment of the University of Hawaii, and the Oklahoma’s Sarkeys Foundation Professorship III. This research is a product of the course “PEPS/MBBE 627 Molecular Diagnostics: Principles and Practices.” The mention of specific trade names or commercial products in this publication does not constitute an official endorsement or recommendation by the University of Hawaii or the USDA.

